# Towards accurate high-throughput ligand affinity prediction by exploiting structural ensembles, docking metrics and ligand similarity

**DOI:** 10.1101/574517

**Authors:** Melanie Schneider, Jean-Luc Pons, William Bourguet, Gilles Labesse

**Affiliations:** Centre de Biochimie Structurale, CNRS, INSERM, Univ Montpellier, 34090 Montpellier, France.

## Abstract

**Motivation:** Nowadays, virtual screening (VS) plays a major role in the process of drug development. Nonetheless, an accurate estimation of binding affinities, which is crucial at all stages, is not trivial and may require target-specific fine-tuning. Furthermore, drug design also requires improved predictions for putative secondary targets among which is Estrogen Receptor alpha (ER*α*).

**Results:** VS based on combinations of Structure-Based VS (SBVS) and Ligand-Based VS (LBVS) is gaining momentum to help characterizing secondary targets of xenobiotics (including drugs and pollutants). In this study, we propose an integrated approach using ligand docking based on multiple structural ensembles to reflect the conformational flexibility of the receptor. Then, we investigate the impact of the two different types of features (structure-based docking descriptors and ligand-based molecular descriptors) for affinity predictions based on a random forest algorithm. We find that ligand-based features have limited predictive power (*r*_*P*_ =0.69, *R*^2^=0.47), compared to structure-based features (*r*_*P*_ =0.78, *R*^2^=0.60) while their combination maintains the overall accuracy (*r*_*P*_ =0.77, *R*^2^=0.56). Extending the training dataset to include xenobiotics, leads to a novel high-throughput affinity prediction method for ER*α* ligands (*r*_*P*_ =0.85, *R*^2^=0.71). Method’s robustness is tested on several ligand databases and performances are compared with existing rescoring procedures. The presented prediction tool is provided to the community as a dedicated satellite of the @TOME server.

**Availability:** http://atome4.cbs.cnrs.fr/ATOME_V3/SERVER/EDMon_v3.html

## 1 Introduction

Despite the fact that the efforts invested in drug development have constantly increased during the last decades, the number of drug approvals stays almost constant (Munos, 2009). Indeed, about 81% of all new drug candidates fail (DiMasi *et al.*, 2010), mainly due to a lack of drug efficiency and/or side effects associated with off-target binding. In order to reduce time and cost of drug development process, various computer aided methods have been implemented. Two main techniques, namely Structure-based and ligand-based virtual screening, are widely used (Lionta *et al.*, 2014; Lavecchia, 2015). They are now used in routine for hit identification in order to prioritize compounds for experimental assays and they are also gaining interest for lead optimization.

**Ligand-based virtual screening** (LBVS) methods are based on analyzing features of substructures and chemical properties related to activity of the ligand. They are useful to search chemical libraries using global or substructure similarity (Mestres and Knegtel, 2000), shape-matching (Nicholls *et al.*, 2010) or pharmacophores (Yang, 2010). The algorithms used in those methods are in constant development and recent LBVS methods are based on data mining and machine learning (Lavecchia, 2015). They do not require structural knowledge, but instead large datasets of characterized ligands.

**Structure-based virtual screening** (SBVS) can be used to predict the binding mode of drugs, to define the important specific interactions between ligand and target and finally to discover a way to improve a given drug by guiding further optimization. SBVS includes docking of candidate ligands into a protein target, followed by evaluation of the likelihood of binding in this pose using a scoring function with an important trade-off between speed and accuracy (Cerqueira *et al.*, 2015). Compared to LBVS, which is restricted to similar molecules the training had been performed on, SBVS is applicable to completely new molecules but it requires knowledge of the targeted structure (or reliable theoretical models). Moreover, affinity cliffs caused by steric clashes, which result from very small changes of the molecules, are more likely to be identified by SBVS methods than by LBVS.

**Combining LBVS with SBVS** is emerging as a way to compensate limitations of each of these complementary approaches. Indeed, there are new attempts to combine both, thanks to the increasing number of both atomic structures and affinity measurements. Usually, the combination of LBVS and SBVS is performed in a sequential or parallel manner (Yu *et al.*, 2018; Zhang *et al.*, 2017). The sequential approach uses both methods as filter steps in a hierarchical procedure with increasing refinement. The parallel approach compares the selected compounds of both methods and retrieves either a consensus (selected by both) or a complementary selection (top molecules from each approach) (Lavecchia and Di Giovanni, 2013). In recent works, sequential selection used interaction fingerprint similarity on large docking outputs. Alternatively, one might apply a weak similarity restraint such as a molecular shape restraint for the ligand (to be classified as a shape-matching LBVS method) during the docking process in SBVS as it is implemented in the docking software PLANTS (Korb *et al.*, 2009).

In the present study, we take advantage of a new interface between PLANTS and the web server @TOME (Pons and Labesse, 2009) to screen multiple conformations in parallel (to be described in more details elsewhere). It also allows us to systematically deduce shape restraints and binding site boundaries based on the geometry of the original ligand from the crystal structure in a fully automatic manner. Subsequent postprocessing is performed using various chemoinformatics tools including several scoring functions to predict protein-ligand affinity.

Ultimately, all the parameters computed to evaluate a ligand pose can be used for machine learning. Indeed, the combination of LBVS and SBVS with machine learning is an emerging approach to improve affinity prediction (Wójcikowski *et al.*, 2017). Therefore, we evaluate applicability of machine learning on the docking outputs of @TOME and PLANTS and ligand similarity measurements. In order to set up and evaluate this development, we focused on a well known therapeutic target - the estrogen receptor ER*α*.

**The ER***α* is a steroid binding receptor playing a key role in a variety of diseases due to its important role in development and physiology. The most prominent examples are ER-based cancer therapies that focus on blocking estrogen action in targeted tissues, with ER*α* being the main target for treatment of ER-positive breast cancer (Ma *et al.*, 2009). The development of new and improved selective ER modulators is therefore still of high interest for pharmaceutical companies to target tissues selectively and to avoid resistance and adverse effects (Wang *et al.*, 2018; Katzenellenbogen *et al.*, 2018; Baker and Lathe, 2018).

On the other hand, ER*α* can also be the unwanted target of drugs or xenobiotics (Delfosse *et al.*, 2012; Baker and Lathe, 2018) and has been identified as anti-target that should be considered in toxicity tests during drug development. Thus, a better understanding of the mechanism of ligand recognition by ER*α* is of paramount importance for safer drug design. Previously, dedicated prediction methods have been addressing the question of whether a molecule is binding or not (Niu *et al.*, 2016; Pinto *et al.*, 2016; Ribay *et al.*, 2016; Mansouri *et al.*, 2016), and traditional structure-activity relationship (QSAR) modeling studies have been also performed with varying success on this nuclear receptor (Waller *et al.*, 1995; Waller, 2004; Asikainen *et al.*, 2004; Zhang *et al.*, 2013; Zhao *et al.*, 2017; Hou *et al.*, 2018).

Despite the fact that ER*α* is an already well characterized therapeutic target (Ekena *et al.*, 1997; Nettles *et al.*, 2004), we are still lacking an efficient and robust method for predicting the binding mode and affinity of docked ligands. A large number of ER*α* crystal structures in complex with ligands are now known and the binding affinity of hundreds of chemical compounds have been experimentally determined. Therefore, ER*α* represents a perfect example to attempt a full characterization by combining SBVS with LBVS and employing machine learning in order to better predict binding affinity and potential future drug profiles.

## 2 Approach

Here, we present an integrated approach for high accuracy affinity predictions on the well known and intensively studied drug target ER*α*. First, a training set was built by systematic docking of chemical compounds extracted from the BindingDB into the available crystal structures of ER*α*. Scoring functions and other chemometric information were gathered for the corresponding complexes and the ligand. Then, we employed a random forest machine learning algorithm on these features ranging from structure-based docking metrics to ligand-based molecular descriptors. All virtual screening results are made available at http://atome4.cbs.cnrs.fr/htbin-post/AT23/MULTI-RUN/FILTER/showform.cgi?WD=AT23/EG/38751543 and the developed prediction tool is provided to the community as an automatic prediction extension within the @TOME-EDMon server (http://atome4.cbs.cnrs.fr/ATOME_V3/SERVER/EDMon_v3.html).

## 3 Materials and methods

### 3.1 Ligand datasets

#### 3.1.1 BindingDB dataset

Two sets of experimentally tested ligands for the human ER*α* (UniProtID: P03372) were extracted from BindingDB (Liu *et al.*, 2007; Gilson *et al.*, 2016) (2018 dataset, updated 2018-04-01). One set contains ligands with known inhibitory constant (Ki) as affinity measure, and a second set contains ligands with half maximal inhibitory concentration (IC50) as an affinity proxy.

A few peptides and a series of boron cluster containing molecules were removed from both datasets, as it was not possible to generate proper 3D conformations or charges for these molecules. The final sets contained 281 ligands (Ki set) and 1641 ligands (IC50 set), respectively. For training, we preferred to focus on the Ki dataset since it corresponds to more direct measurements while the IC50 dataset was used as a larger dataset for method testing.

#### 3.1.2 In-house xenobiotic dataset

The xenobiotic chemical data that was used first as an external testing dataset and afterwards to build an extended training set, is an in-house dataset of 66 ligands with measured affinities for ER*α* (Grimaldi *et al.*, 2015). These extra compounds correspond mostly to bisphenols, halogenated coumpounds, as well as natural macrocycles micmicking estradiol (or phytoestrogens) but harboring distantly related chemical structures.

#### 3.1.3 FDA ER*α* dataset

In order to have a second external validation, we used the Estrogen Receptor targeted dataset from the Endocrine Disruptor Knowledge Base (EDKB) provided by the U.S. Food & Drug Administration (FDA), named here ER-EDKB dataset. The dataset contains 131 ER binders and 101 non-ER binders including natural ligands and xenochemicals that are structurally different from drug-like molecules. For ER binders, the binding affinity measure is reported as a relative binding activity (RBA), which is based on an assay using rat uteri. Those cell-based measurements are influenced by different factors, such as cellular permeability, and are unfortunately not directly comparable with direct Ki measures. Nevertheless, we predicted affinities using all models and transformed the measured RBA values back to pIC50 values (*pIC*50 = *log*_10_(*RBA*) − 8).

### 3.2 Generation of ligand conformations

On the ligand side, there are two factors that can have an impact on docking. One is the initial conformation submitted to a docking program. The second factor are the atomic partial charges that have an impact on ligand pose evaluation (scoring) and can be calculated using different models. The initial ligand sets were downloaded from BindingDB (BDB) and have 3D conformations generated by VConf and partial charges generated by VCharge (Chang and Gilson, 2003). We also tested two other charge models (Gasteiger and MMFF94 charges) instead of the default charge for the 3D conformers built by Vconf. Two other 3D generators (OpenBabel (OLBoyle *et al.*, 2011) and Frog2 (Miteva *et al.*, 2010)) using their default charge (Gasteiger and MMFF94, respectively). This resulted in a total of 5 ligand sets. The ligand sets were then grouped based on variation on their 3D generation, their charges or all together as depicted in Figure 1.

**Figure 1.**
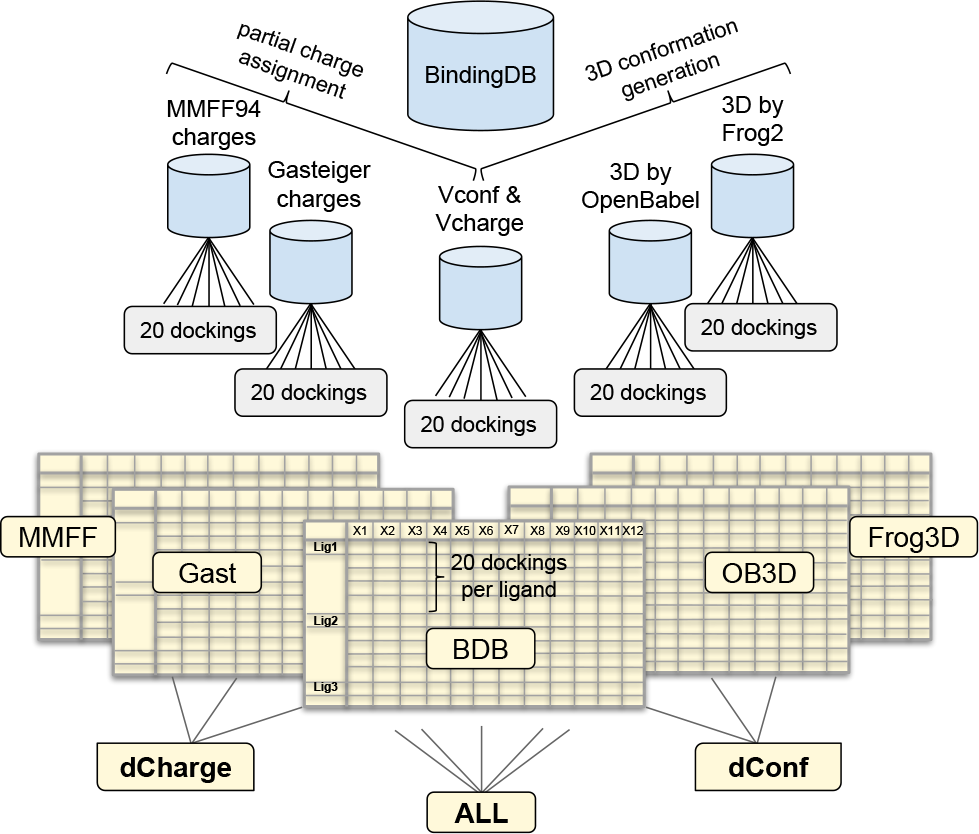
Structure-based dataset generation approach. The ligand dataset was extracted from the BindingDB (BDB), which uses VConf for 3D conformation generation and VCharge for charge assignment. Two more partial charge models (MMFF and Gasteiger) and two other 3D conformation generators (openbabel and FROG-2) were employed to generate a total of 5 ligand sets. Those were submitted to the @TOME server for docking and complex evaluation. The @TOME output datasets ‘MMFF’, ‘Gast’, ‘BDB’, ‘OB3D’ and ‘Frog3D’ (containing the results of 20 dockings per ligand in different structures) were grouped in three combined datasets, a different charge dataset ‘dCharge’, a different conformation dataset ‘dConf’, and an ‘ALL’ dataset.

### 3.3 Structure-based ligand docking

#### 3.3.1 Ensemble docking

First, all liganded ER*α* structures available in the PDB (461 monomers) were gathered using the @TOME server by submitting the ‘canonical’ amino acid sequence of ER*α* (UniProt identifier: P03372-1) with a specified sequence identity threshold of 90%. All gathered 461 monomers had a sequence identity between 95% and 100% with the submitted sequence and correspond to point mutants of the human ER*α*. Missing or substituted side-chains were modeled using SCWRL 3.0 (Wang *et al.*, 2008) using the strictly conserved side-chains fixed. By default, for each ligand to be docked (e.g. from BDB), a set of 20 different template structures were automatically selected among all available PDB structures. This selection is based on the highest similarity (Tanimoto score) between the uploaded ligand and the co-crystallized ligand present in a template. The automatic virtual screening procedure implemented in the @TOME server uses the docking program PLANTS with its shape restraint functionality (with a weighting of −3), using the original ligand of the screened structure as a pharmacophore. Of note, this ligand is also used to define the boundaries of the binding site to be screened (using a distance cutoff of 8 Å). So, not only the protein conformation is (slightly) distinct but various cavity volume and extent are used in this parallel docking procedure. For each template screened, only the best pose was kept. After docking and structure alignment, the 20 computed poses were clustered by conformation similarity, and the most likely pose is selected automatically among the largest cluster using a dedicated heuristics. Accordingly, we perform ligand docking on conformational ensemble as an optimal procedure for SBVS.

#### 3.3.2 Structure-based molecular descriptors

Each docking pose is evaluated by various chemoinformatics tools (see Table 1. Here, we take advantage of several re-scoring functions (namely MedusaScore (Yin *et al.*, 2008), DSX (Neudert and Klebe, 2011), or X-SCORE (Wang *et al.*, 2002)) recently embedded in @TOME to derive a consensus score (including also ChemPLP as used in PLANTS (Korb *et al.*, 2006)) inorder to predict protein-ligand affinities. In addition, other evaluations are performed including model quality (e.g.: Qmean) and internal ligand conformation.

**Table 1.** Structure-based docking metrics

| Metric name | Short description |
| --- | --- |
| PlantsFull | PLANTS score (with anchor weight) |
| Plants | PLANTS ChemPLP score (without weight) |
| PlantsLR | PLANTS pKa (calculated by linear regression on PDBbind) |
| MedusaScore | Medusa original score |
| MedusaLR | MedusaScore pKa (calculated by linear regression on PDBbind) |
| XScore | XScore affinity score (pKa) |
| DSX | DSX original score |
| DSXLR | DSX pKa (calculated by linear regression on PDBbind) |
| AtomeScore | @TOME pKa = mean(PLANTS, XScore, MedusaScore, DSX) |
| Tanimoto | Similarity between candidate ligand and anchor ligand |
| AtomSA | S.A. @TOME score |
| QMean | QMean score of receptor model |
| AnchKd | Affinity calculated between receptor/anchor (pKa) |
| AnchorFit | Candidate/ligand superimposition score (PLANTS software) |
| LigandEnergy | Internal energy of ligand (AMMP force field) |
| LPC | LPC software score (receptor/ligand complementarity function) |
| PSim | Similarity to receptor/ligand interaction profile in PDB template |
| CpxQuality | Complex quality consensus score |
| LPE | Ligand Position Error (SVM multi-variable linear regression) |

The above parameters were important for structure-based screening, and they were complemented by other information regarding the chemical nature of ligands using additional molecular descriptors.

#### 3.3.3 Ligand molecular descriptors

In order to include more information about the small molecules being screened, molecular descriptors were calculated using the Chemistry Development Kit (CDK), a collection of open source Java libraries for chemoinformatics (Guha, 2007). The descriptors were selected based on their ability to represent the diversity of the ligand dataset, taking into account their orthogonality, and based on their variable importance score during model training. The final set of 11 QSAR molecular descriptors includes topological, geometrical, constitutional and charge based descriptors and is listed in Table 2 with descriptor name and a short description.

**Table 2.** Ligand-based molecular descriptors

| Descriptor name | Short description |
| --- | --- |
| Weight | molecular weight |
| VABC | volume descriptor |
| AtomCount | number of atoms |
| BondCount | number of bonds |
| RotatableBondsCount | number of rotatable bonds |
| AromaticBondsCount | number of aromatic bonds |
| HBondDonorCount | number of hydrogen bond donors |
| HBondAcceptorCount | number of hydrogen bond acceptors |
| TPSA | Topological Polar Surface Area |
| XLogP | prediction of logP based on the atom-type method called XLogP |
| HybridizationRatio | fraction of sp3 carbons to sp2 carbons |

#### 3.3.4 Combined structure/ligand descriptors

All 5 docking datasets (originating from the 5 different ligand sets) provided 19 structure-based docking metrics for the 20 docking poses computed for each ligand. For each metric, median and standard deviation were computed and used as a unified instance. Ligand-based variables (11 CDK molecular descriptors) were added to the 19 structure-based metrics. A correlation matrix with all descriptors used for the Ki-BDB dataset is provided as heatmap (see Figure S3). Alternatively, the commonly used MACCS fingerprints (166 features) were also tested for comparison.

### 3.4 Machine learning approaches

#### 3.4.1 Algorithm selection and training

For all analyses, calculations and machine learning, the R language (version 3.2.4) with RStudio was used. The random forest (RF) algorithm provided by the R package ‘randomForest’ showed the highest accuracy in an initial test (see Figure S1) when compared to 6 other machine learning algorithms: linear regression (LinReg), decision tree (CARTree), Gradient Boosting Machine (GBM), support vector machine (SVM) with a linear kernel (SVM_L), a polynomial kernel (SVM_P), and a radial kernel (SVM_R).

All algorithms were employed with default variable settings and 10-fold cross validation. Major advantages of the RF algorithm are that it handles non-linearities, numerical and categorical variables, and it gives estimates of variable importance and generalization error. Therefore, random forest was chosen as the most promising algorithm, since it seemed the best adapted to our data. In order to avoid over-fitting of the models we used stratified 10-fold cross validation repeated 10 times for all models (unless otherwise indicated).

Alternatively, an external test set was built by taking a stratified selection of 20% of the whole dataset. The remaining 80% was used as training set for the models.

#### 3.4.2 Comparison of different tree-based algorithms

The RF algorithm we used, has only one tunable hyperparameter that can be adjusted for the present dataset. Therefore, we wondered whether other tree-based ensemble algorithms with more tunable hyperparameters offer an improved prediction accuracy when tuned more carefully. In total, five different tree-based algorithms were employed on the same Ki BindingDB2018 dataset for affinity prediction and subsequent performance comparison. They are: random forest (RF), regularized random forest (rRF), global regularized random forest (rRFglobal), Extreme Gradient Boosted Trees (xgbTree), and Extreme Gradient Boosted Trees with dropout (xgbDART). Here, Bayesian optimization was employed to select the best hyperparameters (5 to 7 depending on the method), which demands a substantially increase in computational expense compared to the one-variable optimization required for the RF algorithm.

### 3.5 Random forest regression modeling

Random forest models were then trained on each dataset separately (‘MMFF’, ‘Gast’, ‘BDB’, ‘OB3D’ and ‘Frog3D’), on the combination of the 3 different 3D conformation datasets ({‘BDB’, ‘OB3D’, ‘Frog3D’} = ‘dConf’), on the combination of the 3 different partial charge datasets ({‘MMFF’, ‘Gast’, ‘BDB’} = ‘dCharge’), and on all 5 datasets combined (‘ALL’) (compare Figure 1).

Besides the Pearson correlation, two further regression evaluation metrics were used to evaluate the model performance on the external test set. First, the coefficient of determination (*R*^2^) is calculated using the sum of squares method. The second metric, the RMSE, also termed as Root Mean Squared Deviation (RMSD) is the average deviation of the predictions (predicted affinities) from the observations (measured affinities).

## 4 Results and Discussion

We developed and tested an automated and integrated structure- and ligand-based approach to predict quickly accurate binding affinities for ER*α*. This approach takes into account structural variability from the ligand side by using different 3D generators and different charge models, and from the receptor side by using 20 structures for each ligand to be docked. Here, we give access to the docking poses while we evaluate thoroughly the affinity predictions performed using various methods.

### 4.1 Predictions using re-scoring methods

In a first attempt, the predictive power of the four different scoring functions implemented in the @TOME server was assessed.

The Pearson correlations between affinity measurement and the median scores (calculated on 20 docking poses per ligand) are very low for all generated datasets (see Table 3). Even the most recent scoring functions (MedusaScore and DSX) performed poorly in this test. Interestingly, the selection of the best pose among the 20 computed ones slightly improves the correlation between predicted and measured affinities for 3 scoring functions but for MedusaScore which appeared as the most robust and the best for the various ligand description schemes.

**Table 3.** Pearson correlations (r_P_) on all five datasets between experimental affinities and scores from four scoring functions Plants, MedusaScore, DSX and XScore, of (1) the best pose selected by @TOME, and of (2) the median scores of the four scoring functions, calculated on 20 dockings per ligand on all five datasets.

| Dataset name | Plants | MedusaScore | DSX | XScore |
| --- | --- | --- | --- | --- |
| (1) | $r_P$ on predictions for the best pose | | | |
| Gast | 0.042 | 0.154 | 0.129 | 0.060 |
| MMFF | 0.063 | 0.182 | 0.157 | 0.082 |
| BDB | 0.038 | 0.111 | 0.118 | 0.076 |
| OB3D | 0.109 | 0.180 | 0.143 | 0.129 |
| Frog3D | 0.022 | 0.132 | 0.118 | 0.040 |
| (2) | $r_P$ on predictions over 20 poses | | | |
| Gast | -0.031 | 0.204 | 0.019 | 0.049 |
| MMFF | -0.025 | 0.192 | 0.038 | 0.054 |
| BDB | -0.019 | 0.087 | 0.022 | 0.059 |
| OB3D | -0.017 | 0.175 | 0.008 | 0.050 |
| Frog3D | -0.048 | 0.199 | 0.005 | 0.036 |

However, the overall correlation is too low for fine ligand ranking and indicates a limitation of the general-purpose scoring functions. This prompted us to develop a more sophisticated method that should be able to combine advantages of different docking evaluations (structure-based and ligand-based ones) and potentially take into account specific features.

### 4.2 Random Forest regression - model training

#### 4.2.1 Structure-based and ligand-based partial models

To investigate the actual affinity prediction capabilities of structure-based and ligand-based variables, partial models were trained using the 19 structure-based metrics or the 11 Ligands-based metrics on the same dataset named BDB. The docking-metrics only model (*r*_*P*_= 0.780 and *R*^2^= 0.596) outperforms the molecular-descriptors only model (*r*_*P*_= 0.689, and *R*^2^= 0.472) while it has a similar Pearson correlation coefficient and *R*^2^ value compared to the combined model (*r*_*P*_= *r*_*P*_ =0.77, *R*^2^=0.56).

#### 4.2.2 Random Forest model trained on MACCS fingerprints

In this context, it might be interesting to add more information regarding the chemical nature of the ligands studied. Instead of using a reduced set of ligand-based parameters, we turned to use a more thorough description based on an extended and popular fingerprints: MACCS. A new random forest model was trained on MACCS fingerprints representing the ligands only, without providing any structural docking data. This resulted in a Pearson correlation of 0.765 and *R*^2^ of 0.571 on the Ki test set, midway between the two partial models compared above (molecular descriptors only model and docking metrics only model). It is outperformed by our final RF model, which combines ligand-based and structure-based information. Combining MACCS with docking-based features slightly improves the overall performance on the training and testing datasets but further evaluation using external datasets (see below) suggested some overfitting.

#### 4.2.3 Structure-based and ligand-based combined models - trained on single and multiple combined datasets

After optimizing model training by parameter tuning, variable selection and engineering, we compared the various models trained on either single datasets (‘MMFF’, ‘Gast’, ‘BDB’, ‘OB3D’ and ‘Frog3D’) or multiple combined datasets (‘dConf’, ‘dCharge’ and ‘ALL’). Whereas the five models trained on single datasets have a *R*^2^ of 0.66 (*±* 0.01), an RMSE of 0.82 (*±* 0.01) and an explained variance of 63.4 (*±* 0.8), the three models trained on multiple datasets have a better *R*^2^ of 0.68 (*±* 0.004), a lower RMSE of 0.78 (*±* 0.008) and an explained variance of 90.6 (*±* 3.7).

The other models that seem to outperform slightly the RF model on the ‘ALL’ dataset are the boosted tree models xgbTree and xgbDART. But the reverse was true when evaluating the corresponding models onto the reference dataset from the FDA (see below). Most of the differences are weak and may not be significant. Accordingly, the more complex implementations did not provide significant increase in performance and they were not studied further.

### 4.3 Random Forest regression - model testing

Most remarkable is the strong increase in accuracy when using either the ‘dConf’ model trained on the 3 different 3D conformation datasets (‘BDB’, ‘OB3D’ and ‘Frog3D’) or the model trained on the total combined ‘ALL’ dataset comprising all 5 datasets (‘MMFF’, ‘Gast’, ‘BDB’, ‘OB3D’ and ‘Frog3D’). Interestingly, using different charge models improves affinity predictions, but slightly less efficiently than using different 3D conformations. This is probably due to the fact that the binding pocket of ER*α* is mostly hydrophobic and therefore the ligands show the same property and partial charges are predominantly found to differ only marginally.

### 4.4 Analysis of variable importance

To assess the impact of the various parameters from structure-based and ligand-based scoring functions, the variable importance was tracked during training of the RF models. The 30 most important variables for the models trained on the ‘ALL’ dataset is shown in Figure 2. Overall, all models have a rather similar variable importance profile (data not shown).

**Figure 2.**
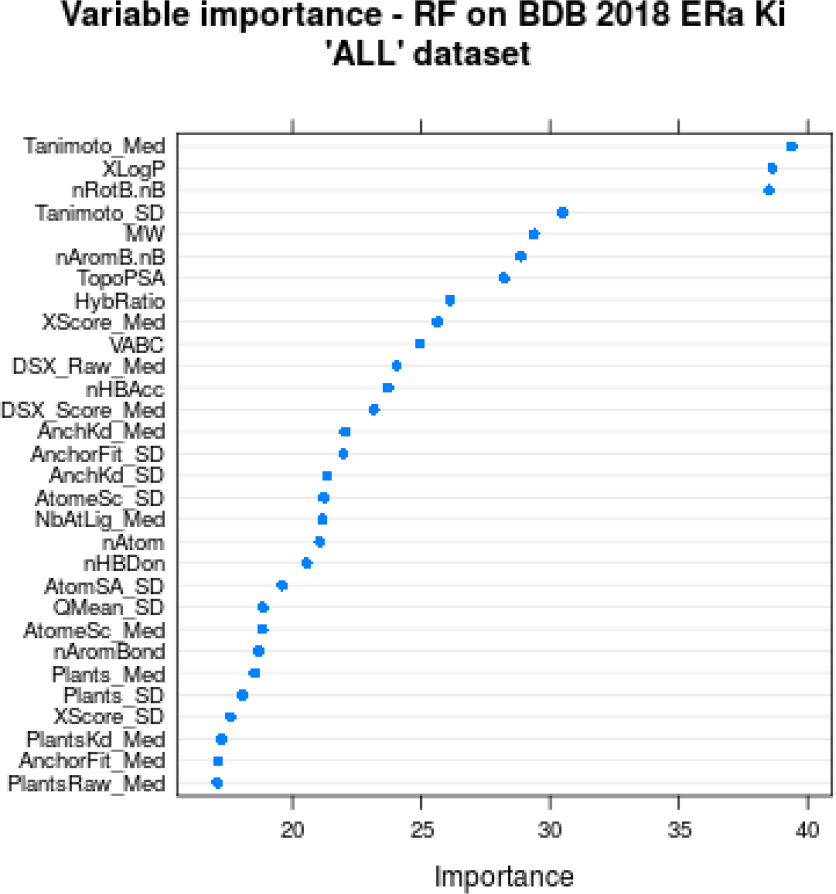
Variable importance of the top 30 variables, tracked during model training for the model trained on the ‘ALL’ dataset with the full variable set. Structure-based docking metrics have an extension (_Med or _SD). The suffix _Med stands for the calculated median of the variable for a ligand’s 20 dockings and _SD is the respective standard deviation of this variable.

**Figure 3.**
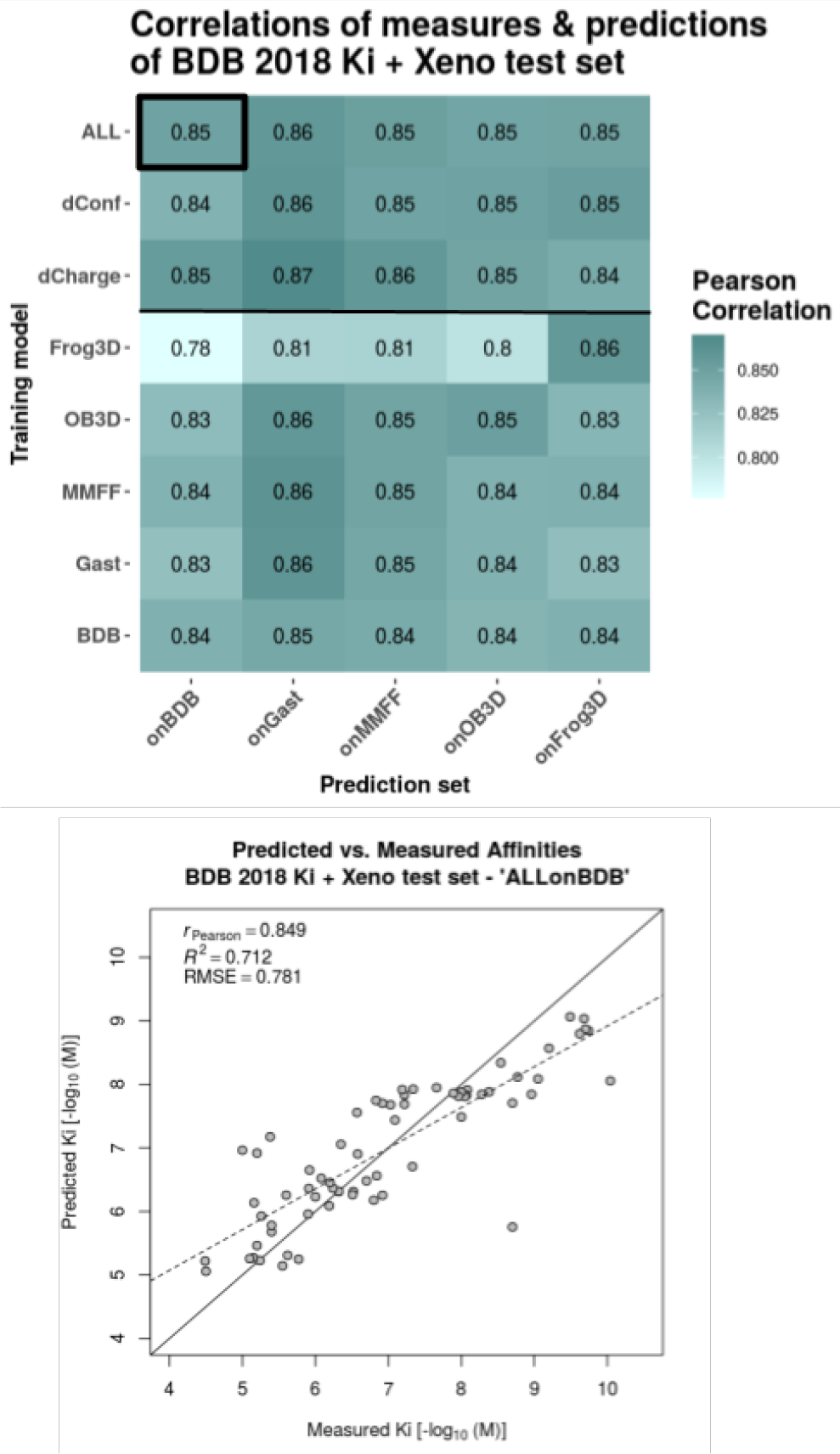
Performance evaluation of extended models on their respective 20% left-out test sets. The initial dataset of 281 ligands is extended by a set of 66 xenochemicals. The heatmap shows Pearson correlations between predictions and measures for all combinations of training model and prediction set. The different training models are listed as rows and the test sets, on which the predictions were made, are listed as columns. RF models were trained on each dataset separately (‘MMFF’, ‘Gast’, ‘BDB’, ‘OB3D’, ‘Frog3D’), on the combination of the 3 different 3D conformation datasets ({‘BDB’, ‘OB3D’, ‘Frog3D’} = ‘dConf’), on the combination of the 3 different partial charge datasets ({‘MMFF’, ‘Gast’, ‘BDB’} = ‘dCharge’), and on all 5 datasets combined (= ‘ALL’). The predictions with the Pearson correlation highlighted in the heatmap (black box) is plotted as scatter-plot for details below. The scatter plot shows the actual predicted versus measured affinities together with a regression line (dashed line), the optimal prediction line (solid diagonal) and the evaluation metrics - Pearson correlation coefficient (rP), coefficient of determination (*R*^2^) and root-mean-square error (RMSE). All evaluation metrics were calculated with respect to the actual values (solid diagonal), not the regression line.

Noteworthy, the most important variable *‘Tanimoto_Med’* is the same for all trained models showing its outstanding importance. It represents the median Tanimoto score calculated between the docked ligand and the 20 shape restraints (or ‘anchors’) present in the targeted structure. This may reflect the importance of using structures bound to similar ligand to ensure proper affinity predictions.

The second and third most important variables are *‘nRotB.nB’* and *‘XLogP’*. *‘nRotB.nB’* estimates ligand flexibility, deduced from the number of rotatable bonds *‘nRotB’* and the total number of bonds *‘nB’* by simply dividing them (*‘nRotB’*/*‘nB’*). During variable testing, this combined variable showed an increased importance compared to the original variables (data not shown), which were therefore removed for the final model training. The particular importance of *‘nRotB.nB’* indicates the important role of entropy cost for binding flexible ligands. Obviously, this parameters is not easily handled in a systematic manner by general scoring functions while it is an important parameter for affinity predictions. In the particular case of ER*α*, it likely discriminates rather small and rigid agonists from larger and more flexible antagonists to prevent overestimating the affinity of the latter. In agreement, the fifth variable is the molecular weight (*‘MW’*) which may also compensate for the additive terms of most scoring functions dedicated to affinity predictions.

Another predominantly important and high-rank variable (second in the ‘ALL’ model and third in the ‘dCharge’ and ‘dConf’ models) is *‘XLogP’*. Representing hydrophobicity and solubility of the ligand, it is expected to be an important factor with respect to the mainly hydrophobic binding pocket of ER*α*. Moreover, *‘XLogP’* may reflect solvent-driven entropic effects that are not easily taken into account by usual scoring functions. Indeed, flexibility and solvation-linked metrics can be regarded as useful for a crude estimate of some entropic effects and counterbalance the enthalpy-oriented affinity prediction approach of usual scoring functions.

Finally, the different scoring functions (DSX, Plants, MedusaScore and X-score; through their means and standard deviations) show a smaller importance than the three above parameters, which could be in agreement with the poor correlations described above. It may also arise from the intrinsic redundancy of our selected variables as several affinity predictionsm are performed in parallel.

Overall, this result underlines the importance of developing dedicated models for each target under investigation, in order to account for some specific features including particular desolvation and flexibility properties.

### 4.5 Model evaluation on different datasets

#### 4.5.1 Model evaluation on a in-house xenobiotic dataset

We took advantage of a complementary and independent dataset - the xenobiotic chemical data of 66 ER*α* binders to evaluate the robustness of our models. Our models performed rather poorly on this dataset with a Pearson correlation of 0.48 for the best Random Forest model BDB-Ki (and 0.40 with the BDB-Ki+MACCS model). Importantly, the chemical nature of most xenobiotics differs significantly from most of the drug-like compounds from the BDB dataset used for training. As such, small xenobiotics (including the small bisphenols) occupy only partially the hydrophobic cavity and often also present numerous halogen substitutions (that are notoriously hard to model). Furthermore, for some of the small xenobiotics we cannot rule out the possibility that two molecules may bind simultaneously (with synergetic effects). This result prompted us to combine these xenobiotics and BDB Ki dataset into an extended training set to build a new RF model.

#### 4.5.2 Model evaluation on FDA ER-EDKB dataset

We then evaluated our two best models on a reference dataset comprising both 322 drug-like and xenobiotic compounds. At the first glance, the predictions made using the original model (trained on only BDB Ki) showed a lower performance especially on the edges of the affinity ranges with both overestimated affinities for small and weak binders (e.g.: alkylphenol) and underestimated predictions for tight binders such as rigid and compact agonists. Indeed, the BDB dataset is mainly composed of large and high-affinity antagonists. Accordingly, some FDA compounds such as high affinity agonists, appear as strong outliers.

Most remarkable is the benefit of adding a complementary dataset of 66 xenobiotic compounds to the initial 281 ligands from BindingDB (see Table 4). Accordingly, the nature and diversity of the ligands matter, so that, proper coverage of the studied chemical space, in the training dataset compared to the testing one is essential.

**Table 4.** Model performances on the FDA ER-EDKB test set. The presented models employ all the RF algorithm and differ in training set composition concerning used molecules and in type of variables used. @TOME+LD = docking evaluation variables from the @TOME server + ligand descriptors calculated with CDK.

| Algorithm | Training set | Variable type | Pearson correlation |
| --- | --- | --- | --- |
| RF | ALL+Xeno | @TOME+LD | 0.748 |
| RF | ALL+Xeno | @TOME+LD+MACCS | 0.740 |
| RF | ALL | @TOME+LD | 0.663 |
| RF | ALL | @TOME+LD+MACCS | 0.648 |
| RF | BDB+Xeno | @TOME+LD | 0.712 |
| RF | BDB+Xeno | @TOME+LD+MACCS | 0.688 |
| RF | BDB | @TOME+LD | 0.584 |
| RF | BDB | @TOME+LD+MACCS | 0.542 |
| RF | BDB+Xeno | MACCS only | 0.487 |

#### 4.5.3 Model evaluation on BindingDB - IC50 dataset

Finally, the most extended and reliable dataset we used for evaluated the our RF models was chosen as the IC50 dataset which includes 1641 entities. Interestingly, the model trained on the Ki dataset only already performed well against IC50 data suggesting a strong robustness.

Training and testing the IC50 dataset (1641 compounds vs 281 for the Ki dataset) provided also some insights into dataset size requirements for the studied target. First, the performance on the IC50 test set (0.87) is better than on the Ki data set (0.77) (compare Table 5). Then, cross-predictions were computed by either using the model constructed on the Ki dataset for predictions on the IC50 dataset, or employing the model constructed on the IC50 dataset for predicting the Ki dataset. In that case, it seems that the small Ki test set (56 compounds) does not allow optimal validation as it shows a significant drop in performance compared to the Ki training set (0.49 vs 0.64). On the contrary, the Ki-ALL model showed similar performance on both the IC50 training and testing sets (1319 versus 322 entities).

**Table 5.** Comparison of cross-predictions between the Ki and IC50 models and datasets. Pearson correlations between experimental affinities and the random forest predictions are reported.

|  | on BDB Ki training set | on BDB Ki test set | on BDB IC50 training set | on BDB IC50 test set |
| --- | --- | --- | --- | --- |
| number of compounds | 225 | 56 | 1319 | 322 |
| Ki ALL model | 0.99 | 0.77 | 0.64 | 0.69 |
| IC50 ALL model | 0.64 | 0.49 | 1.00 | 0.87 |

We also evaluated our last model trained on the extended dataset including both the Ki dataset and the xenobiotic dataset on the largest avaliable IC50 dataset from BindingDB (compare Table 6). Good predictions were observed for the IC50 dataset although the addition of the xenobiotic dataset did not bring any improvement (nor any deterioration) for that particular dataset. Again this suggests that our final model is rather robust.

**Table 6.**
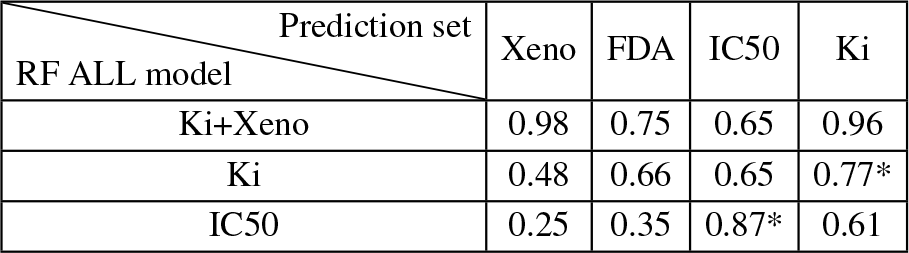
Evaluation of best RF models on various datasets. Pearson correlations between experimental affinities and the RF predictions are reported for the whole datasets but for values marked with ‘*’ that indicates values for a 20% test set.

## 5 Conclusion

We provide an original *in silico* method for accurate binding affinity predictions that takes advantage of structural ensembles, of various structure-based metrics and of ligand-based descriptors in a unique combination. This led to a prediction tool outperforming methods based either solely on SBVS or LBVS approaches as exemplified here with the MACCS fingerprints. Our work confirmed the performance of Random Forest over other machine learning approaches as previously noticed (Russo *et al.*, 2018). In some cases, higher accuracy was reported but for smaller compound libraries (Hou *et al.*, 2018). Accordingly, our results present one of the largest validation surveys and best performing tools for affinity prediction against ER*α*. By training on various types of partial charges and/or 3D builders, we believe our tool will be more robust to variations in the way the submitted compound libraries are generated. Areas for further improvements are probably obtaining increased accuracy in ligand docking, a possible addition of further evaluation metrics for the protein-ligand interactions, as well as using deep learning. Testing challenging compounds is also an important way to guide improvement and we expect our web server to be thoroughly tested with novel compounds.

## Acknowledgements

We thank Muriel Gelin, Corinne Lionne, Dominique Douguet, Matteo Paloni and Rafaela Salgado for careful reading of the manuscript.

## References

Asikainen, A. H., Ruuskanen, J., and Tuppurainen, K. A. (2004). Consensus kNN QSAR: a versatile method for predicting the estrogenic activity of organic compounds in silico. A comparative study with five estrogen receptors and a large, diverse set of ligands. Environmental Science & Technology, 38(24), 6724–6729.

Baker, M. E. and Lathe, R. (2018). The promiscuous estrogen receptor: Evolution of physiological estrogens and response to phytochemicals and endocrine disruptors. The Journal of Steroid Biochemistry and Molecular Biology.

Cerqueira, N. M. F. S. A., Gesto, D., Oliveira, E. F., Santos-Martins, D., Brás, N. F., Sousa, S. F., Fernandes, P. A., and Ramos, M. J. (2015). Receptor-based virtual screening protocol for drug discovery. Archives of Biochemistry and Biophysics, 582, 56–67. 00010.

Chang, C.-E. and Gilson, M. K. (2003). Tork: Conformational analysis method for molecules and complexes. Journal of Computational Chemistry, 24(16), 1987–1998.

Delfosse, V., Grimaldi, M., Pons, J.-L., Boulahtouf, A., le Maire, A., Cavailles, V., Labesse, G., Bourguet, W., and Balaguer, P. (2012). Structural and mechanistic insights into bisphenols action provide guidelines for risk assessment and discovery of bisphenol A substitutes. Proceedings of the National Academy of Sciences of the United States of America, 109(37), 14930–14935. 00075.

DiMasi, J. A., Feldman, L., Seckler, A., and Wilson, A. (2010). Trends in risks associated with new drug development: success rates for investigational drugs. Clinical Pharmacology & Therapeutics, 87(3). 00362.

Ekena, K., Weis, K. E., Katzenellenbogen, J. A., and Katzenellenbogen, B. S. (1997). Different Residues of the Human Estrogen Receptor Are Involved in the Recognition of Structurally Diverse Estrogens and Antiestrogens. Journal of Biological Chemistry, 272(8), 5069–5075.

Gilson, M. K., Liu, T., Baitaluk, M., Nicola, G., Hwang, L., and Chong, J. (2016). BindingDB in 2015: A public database for medicinal chemistry, computational chemistry and systems pharmacology. Nucleic Acids Research, 44(D1), D1045–1053. 00006.

Grimaldi, M., Boulahtouf, A., Delfosse, V., Thouennon, E., Bourguet, W., and Balaguer, P. (2015). Reporter Cell Lines for the Characterization of the Interactions between Human Nuclear Receptors and Endocrine Disruptors. Frontiers in Endocrinology, 6.

Guha, R. (2007). Chemical Informatics Functionality in R. Journal of Statistical Software, 18(1), 1–16.

Hou, T.-Y., Weng, C.-F., and Leong, M. K. (2018). Insight Analysis of Promiscuous Estrogen Receptor -Ligand Binding by a Novel Machine Learning Scheme. Chemical Research in Toxicology, 31(8), 799–813.

Katzenellenbogen, J. A., Mayne, C. G., Katzenellenbogen, B. S., Greene, G. L., and Chandarlapaty, S. (2018). Structural underpinnings of oestrogen receptor mutations in endocrine therapy resistance. Nature Reviews. Cancer, 18(6), 377–388.

Korb, O., Stützle, T., and Exner, T. E. (2006). PLANTS: application of ant colony optimization to structure-based drug design. In Ant Colony Optimization and Swarm Intelligence, pages 247–258. Springer. 00133.

Korb, O., Stützle, T., and Exner, T. E. (2009). Empirical Scoring Functions for Advanced ProteinLigand Docking with PLANTS. Journal of Chemical Information and Modeling, 49(1), 84–96. 00321.

Lavecchia, A. (2015). Machine-learning approaches in drug discovery: methods and applications. Drug Discovery Today, 20(3), 318–331. 00010.

Lavecchia, A. and Di Giovanni, C. (2013). Virtual screening strategies in drug discovery: a critical review. Current Medicinal Chemistry, 20(23), 2839–2860. 00063.

Lionta, E., Spyrou, G., Vassilatis, D. K., and Cournia, Z. (2014). Structure-based virtual screening for drug discovery: principles, applications and recent advances. Current Topics in Medicinal Chemistry, 14(16), 1923–1938. 00018.

Liu, T., Lin, Y., Wen, X., Jorissen, R. N., and Gilson, M. K. (2007). BindingDB: a web-accessible database of experimentally determined protein–ligand binding affinities. Nucleic Acids Research, 35(Database issue), D198–D201. 00773.

Ma, C. X., Sanchez, C. G., and Ellis, M. J. (2009). Predicting endocrine therapy responsiveness in breast cancer. Oncology (Williston Park, N. Y.), 23(2), 133–142.

Mansouri, K., Abdelaziz, A., Rybacka, A., Roncaglioni, A., Tropsha, A., Varnek, A., Zakharov, A., Worth, A., Richard, A. M., Grulke, C. M., Trisciuzzi, D., Fourches, D., Horvath, D., Benfenati, E., Muratov, E., Wedebye, E. B., Grisoni, F., Mangiatordi, G. F., Incisivo, G. M., Hong, H., Ng, H. W., Tetko, I. V., Balabin, I., Kancherla, J., Shen, J., Burton, J., Nicklaus, M., Cassotti, M., Nikolov, N. G., Nicolotti, O., Andersson, P. L., Zang, Q., Politi, R., Beger, R. D., Todeschini, R., Huang, R., Farag, S., Rosenberg, S. A., Slavov, S., Hu, X., and Judson, R. S. (2016). CERAPP: Collaborative Estrogen Receptor Activity Prediction Project. Environmental Health Perspectives, 124(7), 1023–1033.

Mestres, J. and Knegtel, R. M. A. (2000). Similarity versus docking in 3d virtual screening. Perspectives in Drug Discovery and Design, 20(1), 191–207. 00041.

Miteva, M. A., Guyon, F., and Tufféry, P. (2010). Frog2: Efficient 3d conformation ensemble generator for small compounds. Nucleic Acids Research, 38(Web Server issue), W622–W627.

Munos, B. (2009). Lessons from 60 years of pharmaceutical innovation. Nature Reviews Drug Discovery, 8(12), 959–968. 00701.

Nettles, K. W., Sun, J., Radek, J. T., Sheng, S., Rodriguez, A. L., Katzenellenbogen, J. A., Katzenellenbogen, B. S., and Greene, G. L. (2004). Allosteric Control of Ligand Selectivity between Estrogen Receptors and : Implications for Other Nuclear Receptors. Molecular Cell, 13(3), 317–327.

Neudert, G. and Klebe, G. (2011). DSX: a knowledge-based scoring function for the assessment of protein-ligand complexes. Journal of Chemical Information and Modeling, 51(10), 2731–2745. 00085.

Nicholls, A., McGaughey, G. B., Sheridan, R. P., Good, A. C., Warren, G., Mathieu, M., Muchmore, S. W., Brown, S. P., Grant, J. A., Haigh, J. A., Nevins, N., Jain, A. N., and Kelley, B. (2010). Molecular Shape and Medicinal Chemistry: A Perspective. Journal of Medicinal Chemistry, 53(10), 3862–3886. 00165.

Niu, A.-Q., Xie, L.-J., Wang, H., Zhu, B., and Wang, S.-Q. (2016). Prediction of selective estrogen receptor beta agonist using open data and machine learning approach. Drug Design, Development and Therapy, 10, 2323–2331.

OLBoyle, N. M., Banck, M., James, C. A., Morley, C., Vandermeersch, T., and Hutchison, G. R. (2011). Open Babel: An open chemical toolbox. J Cheminf, 3, 33. 00943.

Pinto, C. L., Mansouri, K., Judson, R., and Browne, P. (2016). Prediction of Estrogenic Bioactivity of Environmental Chemical Metabolites. Chemical Research in Toxicology, 29(9), 1410–1427.

Pons, J.-L. and Labesse, G. (2009). @TOME-2: a new pipeline for comparative modeling of protein–ligand complexes. Nucleic Acids Research, 37(suppl 2), W485–W491. 00060.

Ribay, K., Kim, M. T., Wang, W., Pinolini, D., and Zhu, H. (2016). Predictive Modeling of Estrogen Receptor Binding Agents Using Advanced Cheminformatics Tools and Massive Public Data. Frontiers in Environmental Science, 4.

Russo, D. P., Zorn, K. M., Clark, A. M., Zhu, H., and Ekins, S. (2018). Comparing Multiple Machine Learning Algorithms and Metrics for Estrogen Receptor Binding Prediction. Molecular Pharmaceutics.

Waller, C. L. (2004). A comparative QSAR study using CoMFA, HQSAR, and FRED/SKEYS paradigms for estrogen receptor binding affinities of structurally diverse compounds. Journal of Chemical Information and Computer Sciences, 44(2), 758–765.

Waller, C. L., Minor, D. L., and McKinney, J. D. (1995). Using three-dimensional quantitative structure-activity relationships to examine estrogen receptor binding affinities of polychlorinated hydroxybiphenyls. Environmental Health Perspectives, 103(7–8), 702–707.

Wang, L., Guillen, V. S., Sharma, N., Flessa, K., Min, J., Carlson, K. E., Toy, W., Braqi, S., Katzenellenbogen, B. S., Katzenellenbogen, J. A., Chandarlapaty, S., and Sharma, A. (2018). New Class of Selective Estrogen Receptor Degraders (SERDs): Expanding the Toolbox of PROTAC Degrons. ACS Medicinal Chemistry Letters, 9(8), 803–808.

Wang, Q., Canutescu, A. A., and Dunbrack, R. L. (2008). SCWRL and MolIDE: Computer programs for side-chain conformation prediction and homology modeling. Nature protocols, 3(12), 1832–1847. 00082.

Wang, R., Lai, L., and Wang, S. (2002). Further development and validation of empirical scoring functions for structure-based binding affinity prediction. Journal of Computer-Aided Molecular Design, 16(1), 11–26.

Wójcikowski, M., Ballester, P. J., and Siedlecki, P. (2017). Performance of machinelearning scoring functions in structure-based virtual screening. Scientific Reports, 7, 46710.

Yang, S.-Y. (2010). Pharmacophore modeling and applications in drug discovery: challenges and recent advances. Drug Discovery Today, 15(11–12), 444–450.

Yin, S., Biedermannova, L., Vondrasek, J., and Dokholyan, N. V. (2008). MedusaScore: An Accurate Force-Field Based Scoring Function for Virtual Drug Screening. Journal of chemical information and modeling, 48(8), 1656–1662.

Yu, M., Gu, Q., and Xu, J. (2018). Discovering new PI3k inhibitors with a strategy of combining ligand-based and structure-based virtual screening. Journal of Computer-Aided Molecular Design, 32(2), 347–361.

Zhang, L., Sedykh, A., Tripathi, A., Zhu, H., Afantitis, A., Mouchlis, V. D., Melagraki, G., Rusyn, I., and Tropsha, A. (2013). Identification of putative estrogen receptor-mediated endocrine disrupting chemicals using QSAR- and structure-based virtual screening approaches. Toxicology and Applied Pharmacology, 272(1), 67–76.

Zhang, X., Jiang, H., Li, W., Wang, J., and Cheng, M. (2017). Computational Insight into Protein Tyrosine Phosphatase 1b Inhibition: A Case Study of the Combined Ligand- and Structure-Based Approach.

Zhao, Q., Lu, Y., Zhao, Y., Li, R., Luan, F., and Cordeiro, M. N. D. S. (2017). Rational Design of Multi-Target Estrogen Receptors ER and ER by QSAR Approaches. Current Drug Targets, 18(5), 576–591.

