## Supplement for "Towards accurate high-throughput ligand affinity prediction by exploiting structural ensembles, docking metrics and ligand similarity"

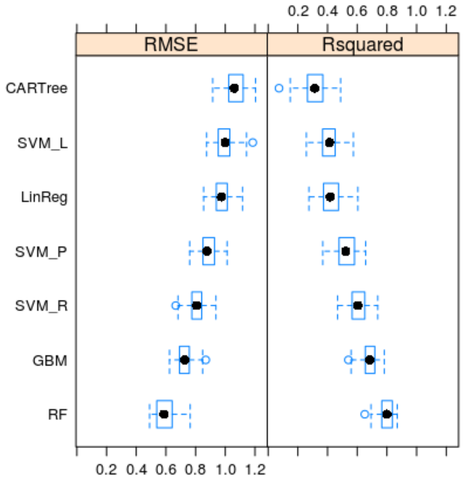


**Figure S1.** Performance comparison of different algorithms for binding affinity prediction (regression) - as boxplot. Models are ranked according to their performance (top = lowest performance, bottom = best performance): Classification and regression tree (CARTree), linear regression (LinReg), support vector machine with linear kernel (SVM_L), with polynomial kernel (SVM_P), and with radial kernel (SVM_R), general boosted machine (GBM), and random forest (RF).

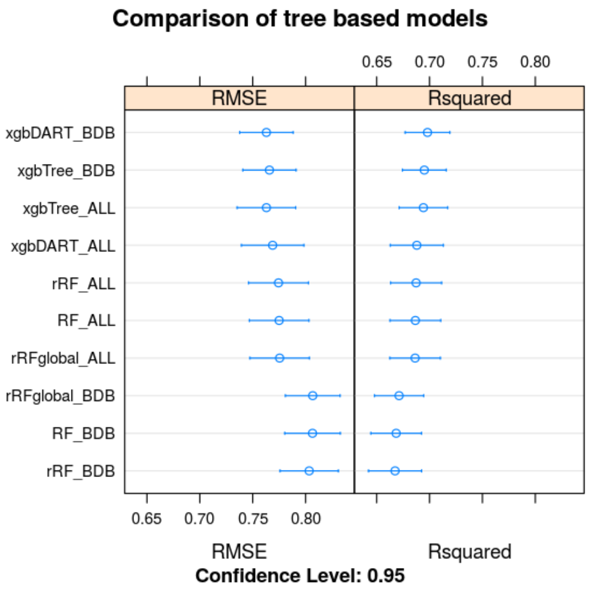


**Figure S2.** Comparison of different tree-based algorithms with performance metrics calculated on cross-validation samples.

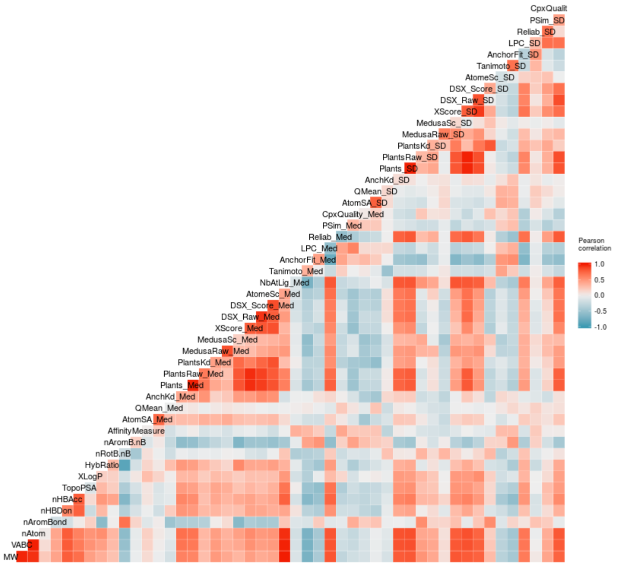


**Figure S3.** Descriptor correlation matrix for the Ki-BDB dataset as heatmap.

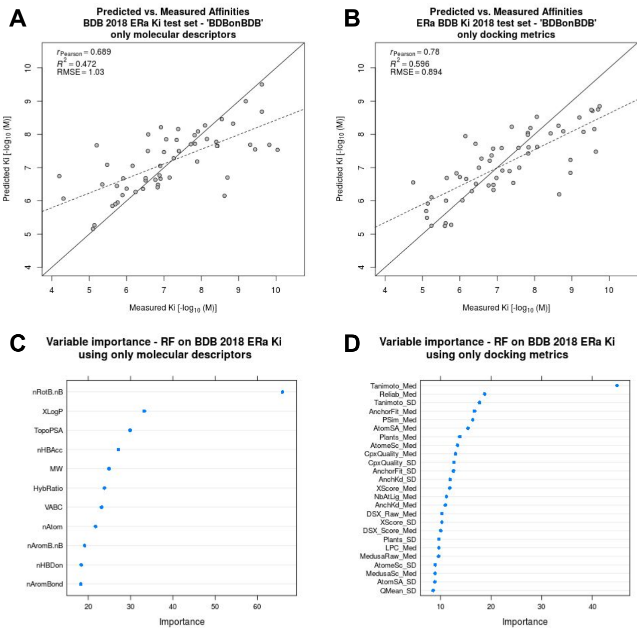


**Figure S4.** Correlations between measured and predicted affinities for the external Ki test set. The predictions were generated by models that were trained on a subset of descriptors. A) model trained only on the set of ligand-based molecular descriptors, and B) model trained only on the set of structure-based metrics from the @TOME server. C) and D) show the ranked variable importance to the trained models A) and B), respectively.

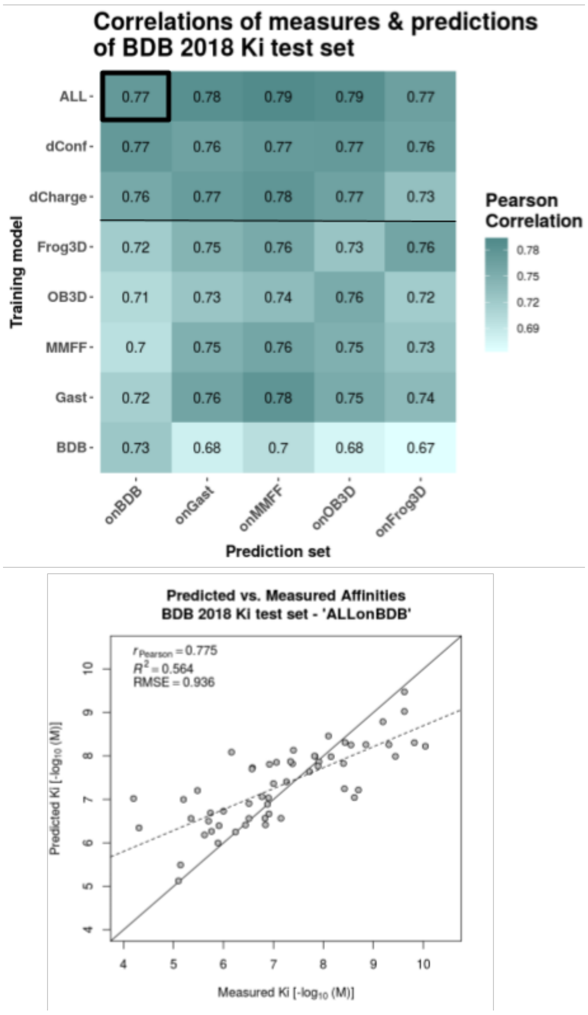


**Figure S5.** Correlations between measured and predicted affinities for the external Ki test set. The heatmap shows Pearson correlations between predictions and measures for all combinations of training model and prediction set. The different training models are listed as rows and the test sets, on which the predictions were made, are listed as columns. Random forest models were trained on each dataset separately ('MMFF', 'Gast', 'BDB', 'OB3D', 'Frog3D'), on the combination of the 3 different 3D conformation datasets ({'BDB', 'OB3D', 'Frog3D'} = 'dConf'), on the combination of the 3 different partial charge datasets ({'MMFF', 'Gast', 'BDB'} = 'dCharge'), and on all 5 datasets combined (= 'ALL'). For one prediction set the scatter plot below shows the actual predicted versus measured affinities together with a regression line (dashed line), the optimal prediction line (solid diagonal) and the evaluation metrics - Pearson correlation coefficient ($r_P$), coefficient of determination ($R^2$) and root-mean-square error (RMSE). All evaluation metrics were calculated with respect to the actual values (solid diagonal), not the regression line.

Table S1. Model performances on the FDA ER-EDKB test set. The presented models differ in algorithm usage, amount of cross-validation folds (by default 10-fold unless indicated differently as 3-CV), and training set composition concerning used molecules. The type of variables used remains unchanged. @TOME+LD = docking evaluation variables from the @TOME server + ligand descriptors calculated with CDK.

| Algorithm | Training set | Variable type | Pearson correlation |
|---|---|---|---|
| xgbTree | ALL | @TOME+LD | 0.677 |
| rRF | ALL | @TOME+LD | 0.670 |
| RF | ALL | @TOME+LD | 0.663 |
| RF(3-CV) | ALL | @TOME+LD | 0.661 |
| rRFglobal | ALL | @TOME+LD | 0.661 |
| xgbDART | ALL | @TOME+LD | 0.618 |
| RF(3-CV) | BDB | @TOME+LD | 0.592 |
| RF | BDB | @TOME+LD | 0.584 |
| rRFglobal | BDB | @TOME+LD | 0.580 |
| xgbDART | BDB | @TOME+LD | 0.540 |
| rRF | BDB | @TOME+LD | 0.519 |
| xgbTree | BDB | @TOME+LD | 0.500 |
